# PHGDH is a targetable driver of PDAC progression

**DOI:** 10.64898/2026.03.11.711147

**Authors:** Yumi Kim, Le Jin Sun, Meredith Long, Samantha Caldwell, H. Carlo Maurer, Kenneth P. Olive, Florian Karreth, Gina M. DeNicola

**Affiliations:** Department of Metabolism and Physiology, H. Lee Moffitt Cancer Center and Research Institute, Tampa, FL 33612, USA; TUM School of Medicine and Health, Clinical Department of Internal Medicine II, TUM University Hospital, Munich, Germany; Department of Medicine, Vagelos College of Physicians and Surgeons, Columbia University Irving Medical Center, New York, NY; Herbert Irving Comprehensive Cancer Center, Columbia University Irving Medical Center, New York, NY; Department of Molecular Oncology, H. Lee Moffitt Cancer Center and Research Institute, Tampa, FL 33612, USA

**Author notes:** **Corresponding Author:** Gina M. DeNicola. **Conflict of Interest:** The authors declare no potential conflicts of interest.

## Abstract

Pancreatic ductal adenocarcinoma (PDAC) arises in a nutrient-deprived microenvironment through progressive stages from pancreatic intraepithelial neoplasia (PanIN) to invasive carcinoma. While serine metabolism supports tumor growth across multiple cancer types, the stage-specific role of de novo serine synthesis in PDAC evolution remains undefined. Here, we show that expression of phosphoglycerate dehydrogenase (PHGDH), the rate-limiting enzyme of serine biosynthesis, increases progressively from PanIN to invasive PDAC in human and mouse specimens. Using genetically engineered mouse models with inducible PHGDH knockdown, we found that PHGDH loss delayed PDAC development. Unexpectedly, PHGDH-deficient tumors did not increase reliance on exogenous serine, and dietary serine/glycine manipulation had no effect on tumor development. Instead, stable isotope tracing and metabolomic profiling revealed that PHGDH loss suppressed mTOR signaling, reduced expression of the glutamine transporter ASCT2, and impaired glutamine uptake and utilization. Leveraging this metabolic liability, we demonstrated that PHGDH-deficient tumors exhibited selective sensitivity to the glutamine antagonist DRP-104, whereas PHGDH-intact tumors were resistant. These findings reveal an unanticipated connection between serine biosynthesis and glutamine metabolism in PDAC and identify a therapeutic vulnerability that may be exploited through combined metabolic targeting.

**Statement of significance:** PHGDH supports PDAC progression not primarily through serine provision, but by maintaining glutamine metabolism and mTOR signaling. This unanticipated metabolic crosstalk creates a synthetic lethal vulnerability to glutamine antagonism in PHGDH-deficient tumors, providing a rationale for combining serine synthesis pathway inhibitors with glutamine-targeting therapies in pancreatic cancer.

## Introduction

Pancreatic ductal adenocarcinoma (PDAC) is one of the most lethal solid malignancies, with high mortality and limited therapeutic options (1, 2). PDAC progression is accompanied by extensive metabolic rewiring that supports tumor growth and survival, highlighting the importance of defining stage-specific metabolic requirements across precursor lesions and malignant transition (3, 4).

PDAC development follows a well-characterized histological progression through acinar-to-ductal metaplasia (ADM) (5, 6) and pancreatic intraepithelial neoplasia (PanIN) stages before the development of invasive carcinoma (7, 8). This progression is accompanied by distinct metabolic rewiring that supports the energetic and biosynthetic demands of transformation. Among the metabolic adaptations observed in PDAC, alterations in serine metabolism present a particularly intriguing paradox. Serine is a central metabolic hub that supports nucleotide synthesis, redox homeostasis, one-carbon metabolism, and lipid biosynthesis (9–13). Serine can be obtained from the diet, *de novo* synthesis, or recycling of intracellular and extracellular components. Dietary restriction of serine and glycine suppresses tumor growth in several murine cancer models (11, 14). Notably, Kras-driven pancreatic cancer is unresponsive to dietary serine and glycine restriction (14). More broadly, serine availability can engage metabolic stress-adaptation programs that shape cell survival and metabolism (12), and increased dependence on de novo serine biosynthesis has been linked to tumor growth and therapeutic resistance across multiple cancer contexts (15, 16). Moreover, while macropinocytosis supports most amino acid pools in PDAC, it does not highly contribute to serine and glycine (17). While other exogenous sources, including neuronal serine release within the PDAC microenvironment (18), may contribute to resistance to dietary restriction, these observations indicate that intrinsic metabolic adaptations could also contribute to dietary serine independence in PDAC.

Upregulation of de novo serine synthesis through the serine synthesis pathway (SSP) represents one such intrinsic adaptation. The SSP converts the glycolytic intermediate 3-phosphoglycerate into serine. Phosphoglycerate dehydrogenase (PHGDH), the first and rate-limiting enzyme of the SSP, supports nucleotide production, redox control, lipid homeostasis, epigenetic regulation, and stress adaptation (9, 10, 19–21). Increased activity of the SSP has been shown in multiple cancers (22–25), driven by genomic amplification (26, 27), transcriptional regulation downstream of MYC (28), and BRAF and KEAP1 mutations (25, 29), and other mechanisms (30). Genetic and functional studies implicate PHGDH as a metabolic driver in cancer, including the requirement for PHGDH for primary melanoma formation (29) and its metastatic survival in serine-low sites such as the brain (31). PHGDH upregulation has also been reported in therapy-resistant contexts, consistent with metabolic adaptation to treatment-induced stress (32, 33). These observations have motivated the development of PHGDH inhibitors, although none has yet demonstrated clear clinical benefit (23, 24). Whether the SSP underlies the intrinsic serine independence of PDAC, and whether this represents a therapeutically targetable dependency, remains unknown.

Here, we delineate the stage-specific role of PHGDH during PDAC progression in vivo. We found that SSP enzymes increase from precursor lesions to invasive PDAC and that PHGDH loss delays progression to carcinoma while promoting apoptosis in established tumors. Unexpectedly, despite reduced serine biosynthesis, PHGDH-deficient tumors did not increase reliance on exogenous serine. Instead, PHGDH loss impaired glutamine handling and created a selective vulnerability to glutamine antagonism, revealing an unanticipated metabolic dependency that may offer therapeutic opportunities.

## Results

### PHGDH expression increases during PDAC progression and contributes to PDAC development

To determine whether the serine synthesis pathway is upregulated during PDAC progression, we profiled laser capture microdissected ductal epithelial cells from human PanIN and PDAC specimens (34). PDAC samples (N = 197) showed higher *PHGDH* and *PSAT1* mRNA expression than PanIN samples (N = 26) (Fig. 1A). A similar pattern was observed in mice, where PHGDH expression was higher in PDAC from KPC mice (LSL-Kras^G12D^; p53^flox/flox^; p48-Cre) than in PanIN from KC mice (LSL-Kras^G12D^; p48-Cre) (Fig. 1B). These findings indicate that SSP expression increases during progression from PanIN to invasive PDAC, raising the question of whether PHGDH is required for progression.

**Figure 1.**
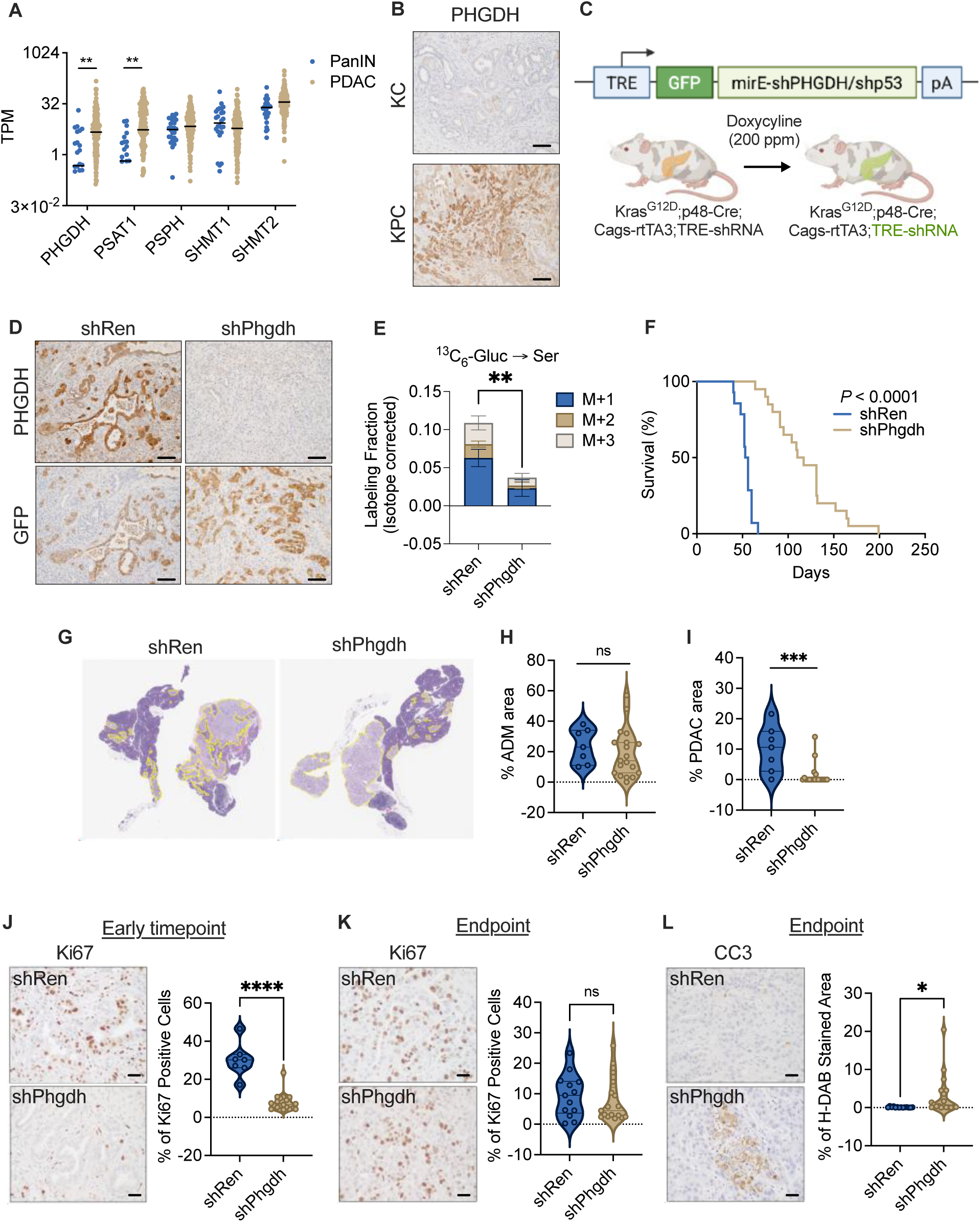
PHGDH upregulation during PDAC progression supports tumor development. (A) Expression of serine synthesis pathway genes in laser capture microdissection–enriched epithelial compartments from human PanIN (n = 26) and PDAC (n = 197) samples. Each dot represents one sample and values are shown as transcripts per million (TPM). (B) Representative PHGDH immunohistochemistry (IHC) of pancreata from LSL-Kras^G12D/+^; p48-Cre (KC) mice collected at 120 days and LSL-Kras^G12D/+^; Trp53^flox/flox^; p48-Cre (KPC) mice collected at 60 days. Scale bars = 50 μm. (C) Schematic of the ESC-GEMM PDAC model with doxycycline-inducible miRE shRNAs. Doxycycline chow (200 ppm) was initiated at postnatal day 30 to induce shp53 together with either control shRen or shPhgdh. GFP serves as a reporter of shRNA induction. (D) Representative IHC staining for PHGDH and GFP in PDAC tissues from shRen and shPhgdh mice collected at endpoint. Scale bars = 50 μm. (E) De novo serine synthesis assessed by infusion of U-^13^C_6_-glucose (150 mg/mL; 3 μL/min for 2.5 hours) via jugular catheterization. Isotopologue-corrected serine labeling (M+1, M+2, M+3) was quantified by LC-MS in shRen and shPhgdh tumors (n = 5 per group). Bars show mean ± SD. **P < 0.01. (F) Kaplan-Meier overall survival curves for shRen (n = 14) and shPhgdh (n = 20) mice. P value was determined by the log-rank (Mantel–Cox) test. (G-I) Representative whole-slide H&E images of pancreata collected 35 days after doxycycline induction from shRen (n = 7) and shPhgdh (n = 19) mice. (G) Yellow outlines indicate regions annotated as disease (including ADM, PanIN, and PDAC). (H) Quantification of ADM area expressed as a percentage of total pancreatic area. (I) Quantification of PDAC area expressed as a percentage of total annotated disease area. (J) Representative Ki67 IHC and quantification of Ki67-positive cells within disease regions in pancreata from shRen (n = 7) and shPhgdh (n = 19) mice collected 35 days after doxycycline induction. Scale bars = 20 μm. (K) Representative Ki67 IHC and quantification of Ki67-positive cells in PDAC tissues from shRen (n = 13) and shPhgdh (n = 17) mice collected at endpoint. Scale bars = 20 μm. (L) Representative cleaved caspase-3 (CC3) IHC and quantification of CC3-positive staining in PDAC tissues from shRen (n = 14) and shPhgdh (n = 19) mice collected endpoint, reported as percentage of H-DAB-positive area. Scale bars = 20 μm. Violin plots show the distribution of values; each dot represents an individual mouse. Center lines indicate the median and interquartile range. P values: *P < 0.05; **P < 0.01; ***P < 0.001; ns, not significant.

To test whether PHGDH is functionally required for PDAC development *in vivo*, we used an embryonic stem cell-genetically engineered mouse model (ESC-GEMM) with pancreas-specific *Kras^G12D^* activation (LSL-Kras^G12D^; p48-Cre) and doxycycline-inducible, GFP-linked shRNAs (35). Starting at postnatal day 30, doxycycline administration induced shRNA expression to silence *Trp53* (shp53) together with either a control shRNA targeting *Renilla luciferase* (shRen) or an shRNA targeting *Phgdh* (shPhgdh), enabling temporally controlled silencing during progression to adenocarcinoma (Fig. 1C). IHC confirmed reduced PHGDH expression in GFP-positive shPhgdh PDAC cells, indicating efficient knockdown (Fig. 1D). Following U-^13^C_6_-glucose infusion, LC–MS analysis showed decreased serine labeling in shPhgdh tumors, consistent with impaired *de novo* serine synthesis (Fig. 1E). PHGDH knockdown prolonged PDAC-free survival compared with shRen controls (Fig. 1F), indicating that PHGDH supports efficient PDAC formation.

To determine whether PHGDH is required during the early stages of disease progression, we induced shRen or shPhgdh with doxycycline and collected pancreata after 35 days for histopathologic analysis. At this timepoint, disease was primarily acinar-to-ductal metaplasia (ADM). PHGDH knockdown had minimal impact on overall disease burden (% of total tissue area) but reduced the PDAC fraction within diseased regions, indicating impaired progression to invasive carcinoma (Fig. 1G–I). Consistent with this, PHGDH knockdown decreased proliferation in ADM lesions (Fig. 1J). By contrast, endpoint PDAC showed comparable proliferation between groups (Fig. 1K), while PHGDH knockdown increased the number of apoptotic PDAC cells (Fig. 1L). These findings suggest PHGDH is preferentially required during the transition to invasive carcinoma, rather than in sustaining proliferation in established PDAC.

### PHGDH-deficient PDAC does not increase reliance on exogenous serine

Impaired *de novo* serine synthesis can potentially be compensated by increased uptake of exogenous serine, as demonstrated in colorectal cancer models (36). If PHGDH-deficient PDAC tumors similarly increase serine scavenging, dietary serine restriction might be synthetically lethal with PHGDH loss. We therefore sought to examine whether PDAC tumors with impaired *de novo* serine synthesis increase their reliance on exogenous serine. We first established a serine and glycine supplementation strategy by providing C57BL/6J mice with drinking water containing 2% serine and glycine (w/v) for 4 weeks. Metabolic profiling confirmed a marked increase in pancreatic serine and glycine levels, with approximately 7.2-fold and 4-fold elevations, respectively, compared with control water (Fig. 2A, B), indicating effective delivery of exogenous amino acids to the pancreas.

**Figure 2.**
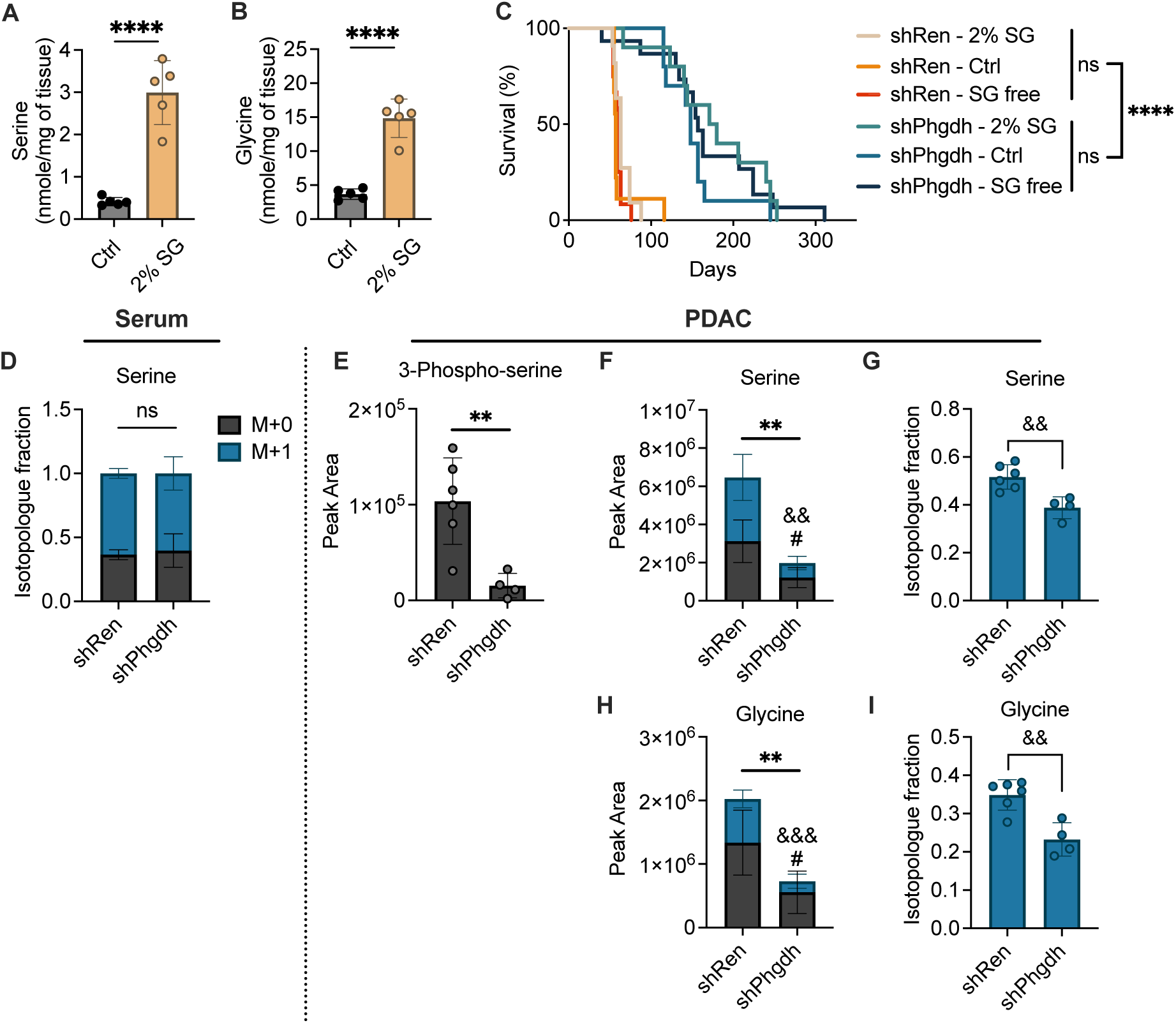
PHGDH knockdown tumors are refractory to serine/glycine dietary manipulation. (A-B) Female C57BL/6NJ mice were provided control drinking water (n = 5) or drinking water supplemented with 2% (w/v) serine and glycine (n = 5) for 28 days. Serine (A) and glycine (B) levels were measured by LC-MS in pancreas tissue. Bars show mean ± SD. ****P < 0.0001. (B) Kaplan-Meier overall survival curves for shRen and shPhgdh PDAC mice maintained on defined diets containing doxycycline (200 ppm) that either included or excluded serine/glycine, with or without 2% (w/v) serine/glycine supplementation in drinking water. Control diet (Ctrl): shRen (n = 9), shPhgdh (n = 10); serine/glycine-free diet (SG-free): shRen (n = 12), shPhgdh (n = 15); Control diet with 2% (w/v) serine/glycine supplementation in drinking water (2% SG): shRen (n = 11), shPhgdh (n = 10). P-values were determined by the log-rank test. ns, not significant; ****P < 0.0001. (D-I) Serine uptake was evaluated by infusion of ^13^C_1_-serine (20 mg/mL) via jugular catheterization at 2 µL/min for 4 hours. ^13^C_1_-serine labeling was quantified by LC-MS in (D) serum and (E-I) PDAC tumors from shRen (n = 6) and shPhgdh (n = 4) mice. Bars represent mean ± SD; each dot represents an individual mouse. Unpaired two-tailed t tests were used for shRen versus shPhgdh comparisons. Symbols denote significance for total pool size (*), M+0 (#), and M+1 (&). *P < 0.05; **P < 0.01; #P < 0.05; &&P < 0.01; &&&P < 0.001; ns, not significant.

Next, we tested whether altering exogenous serine and glycine availability modifies PDAC development in the setting of PHGDH deficiency. Under doxycycline induction, shRen and shPhgdh mice were maintained on a defined amino acid control diet, a serine and glycine-free diet, or a control diet supplemented with either vehicle water or 2% serine and glycine water. Serine and glycine supplementation or deprivation did not affect PDAC-free survival in either genotype, and shPhgdh mice retained a pronounced PDAC-free survival advantage over shRen controls under all conditions (Fig. 2C). These data indicate that tumor onset remains insensitive to serine and glycine availability even when endogenous serine synthesis is genetically impaired. To directly determine whether PHGDH-deficient tumors compensate by increasing exogenous serine uptake and utilization, mice were infused with ^13^C_1_-serine. Serum serine labeling was comparable between shRen and shPhgdh mice, confirming equivalent systemic tracer availability (Fig. 2D). As expected, the SSP intermediate 3-phospho-L-serine was decreased in shPhgdh tumors (Fig. 2E), consistent with reduced SSP activity *in vivo*. In PDAC tissue, PHGDH deficiency did not increase fractional M+1 labeling of serine or glycine relative to shRen controls (Fig. 2F, G). Instead, shPhgdh tumors exhibited lower fractional M+1 labeling of serine and glycine, with decreased total levels of unlabeled (M+0) and labeled (M+1) forms (Fig. 2F-I). These findings indicate that PHGDH loss does not trigger compensatory reliance on exogenous serine during tumor development.

### PHGDH deficiency limits glutamine uptake in PDAC

To define the metabolic consequences of PHGDH silencing in PDAC, we performed global metabolomic profiling of shRen and shPhgdh tumors. This analysis revealed broad metabolic changes in shPhgdh tumors, including reductions in serine, glutamine, threonine, and alpha-ketoglutarate (α-KG) (Fig. 3A), suggesting that PHGDH loss impacts pathways beyond serine production.

**Figure 3.**
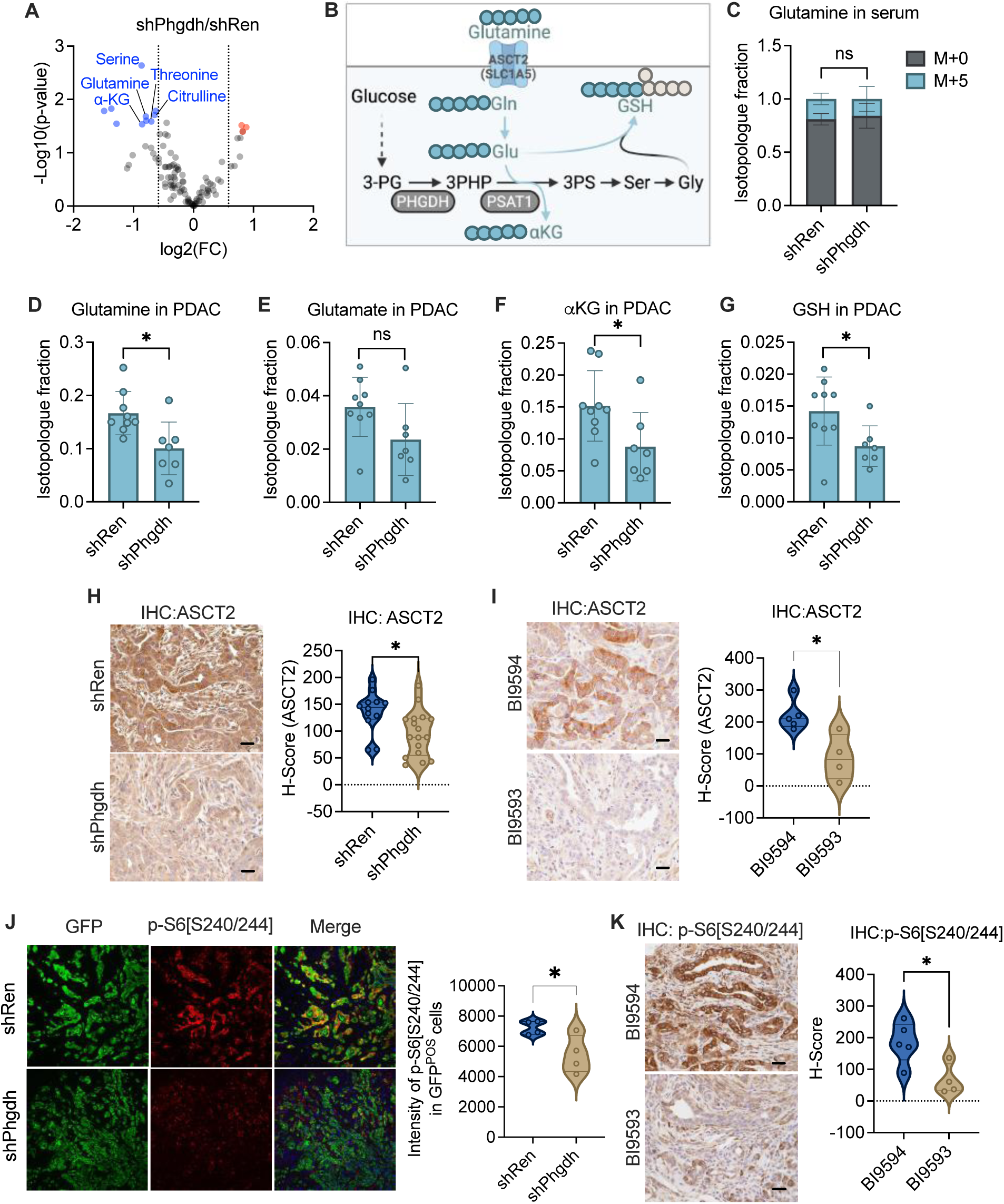
PHGDH deficiency reduces glutamine uptake and the expression of the glutamine transporter ASCT2. (A) Volcano plot of metabolite abundance changes in endpoint PDAC tissues from shRen and shPhgdh mice (n = 9 per group). Selected significantly altered metabolites are annotated. (B) Schematic illustrating the relationship between PHGDH-dependent serine synthesis, ^13^C_5_-glutamine uptake via ASCT2 (SLC1A5), and downstream glutamine-derived metabolites. (C-G) Glutamine uptake and downstream metabolism were evaluated by infusion of U-^13^C_5_-glutamine (15 mg/mL; 2 µL/min) for 4 hours via tail vein catheterization in shRen (n = 9) and shPhgdh (n = 7) PDAC mice. Glutamine isotopologue fractions were quantified by LC-MS: (C) serum glutamine M+0 and M+5, and (D–G) PDAC tumor M+5 fractions for glutamine, glutamate, α-ketoglutarate (αKG), and glutathione (GSH). Bars show mean ± SD; each dot represents one mouse. *P < 0.05; ns, not significant. (H) Representative ASCT2 IHC images and H-score quantification in endpoint PDAC tumors from shRen (n = 12) and shPhgdh (n = 17) mice. Scale bars = 20 μm, *P < 0.05. (I) Representative ASCT2 IHC images and H-score quantification in PDAC tumors from shRen mice treated with BI-9593 (PHGDH inhibitor; n = 4) or BI-9594 (inactive control; n = 5) by oral gavage (250 mg/kg; three doses at 12-hour intervals) prior to tumor collection. Scale bars = 20 μm. Violin plots show distributions with each dot representing an individual mouse. *P < 0.05. (J) Representative immunofluorescence images of GFP (green), p-S6 [S240/244] (red), and Merge (yellow indicates colocalization) in shRen (n = 4) and shPhgdh (n = 4) PDAC tumors collected at endpoint. Violin plots show quantification of p-S6 [S240/244] intensity in GFP-positive regions. *P < 0.05. (K) Representative IHC staining for p-S6 [S240/244] following BI9594 (n = 5) or BI9593 (n = 4) treatment as in (I), with H-score quantification. Each dot represents an individual mouse; Scale bars = 20 μm. * P ≤ 0.05.

Among the depleted metabolites, the reduction in glutamine was particularly striking. Glutamine is a critical nutrient in PDAC, where it supports both anaplerosis and redox balance (37, 38). Moreover, glutamine-derived glutamate serves as the nitrogen donor in the PSAT1-catalyzed transamination reaction during serine biosynthesis (Fig. 3B), creating a potential metabolic link between the SSP and glutamine metabolism. We therefore investigated whether PHGDH loss alters glutamine uptake and downstream glutamine metabolism. Tumor-bearing shRen and shPhgdh mice were infused with ^13^C_5_-glutamine. Serum glutamine M+5 labeling was comparable between groups (Fig. 3C), but tumor glutamine M+5 labeling was significantly reduced in shPhgdh tumors (Fig. 3D), indicating impaired glutamine uptake. Among downstream metabolites, α-KG and glutathione M+5 labeling were significantly reduced in shPhgdh tumors, with glutamate showing the same trend (Fig. 3E-G). Together, these data indicate that PHGDH deficiency impairs glutamine uptake and utilization in PDAC tumors.

We next examined whether PHGDH deficiency alters expression of glutamine transporters. ASCT2 (*Slc1a5*) protein expression was reduced in shPhgdh tumors despite elevated *Slc1a5* mRNA levels (Fig. 3H, Supplementary Fig. 1A, B), indicating post-transcriptional regulation. To determine whether this requires PHGDH enzymatic activity, we treated tumor-bearing mice with the PHGDH inhibitor BI-9593 or inactive control BI-9594. BI-9593 (250 mg/kg) fully suppressed serine biosynthesis, as assessed by U-^13^C_6_-glucose tracing (Supplementary Fig. 2). BI-9593 reduced ASCT2 expression (Fig. 3I), phenocopying genetic PHGDH knockdown.

Additionally, phosphorylated S6 [S240/244] (p-S6), a marker of mTOR-dependent protein synthesis, was reduced in shPhgdh PDAC tumors compared to shRen controls (Fig. 3J). Pharmacologic PHGDH inhibition with BI-9593 similarly reduced p-S6 expression (Fig. 3K), confirming this effect depends on PHGDH enzymatic activity and occurs with acute inhibition rather than as a long-term adaptation to PHGDH loss. These findings demonstrate that PHGDH supports both glutamine metabolism and mTOR signaling in PDAC.

### PHGDH loss creates a selective vulnerability to glutamine antagonism in PDAC

Given that PHGDH deficiency impairs glutamine uptake and utilization, we hypothesized that PHGDH-deficient tumors would be more vulnerable to pharmacologic glutamine antagonism. We therefore tested whether PHGDH silencing alters sensitivity to glutamine antagonism using DRP-104, a pro-drug of the glutamine antagonist DON designed to reduce systemic toxicity and shown to suppress PDAC tumor growth in preclinical models (39). Mice were placed on a doxycycline diet starting at postnatal day 30, and treatment was initiated once tumors reached approximately 50 mm^3^ by ultrasound (Fig. 4A). DRP 104 was administered intraperitoneally at 3 mg/kg on a 5-days-on/2-days-off schedule, and tumor volumes were monitored by serial ultrasound.

**Figure 4.**
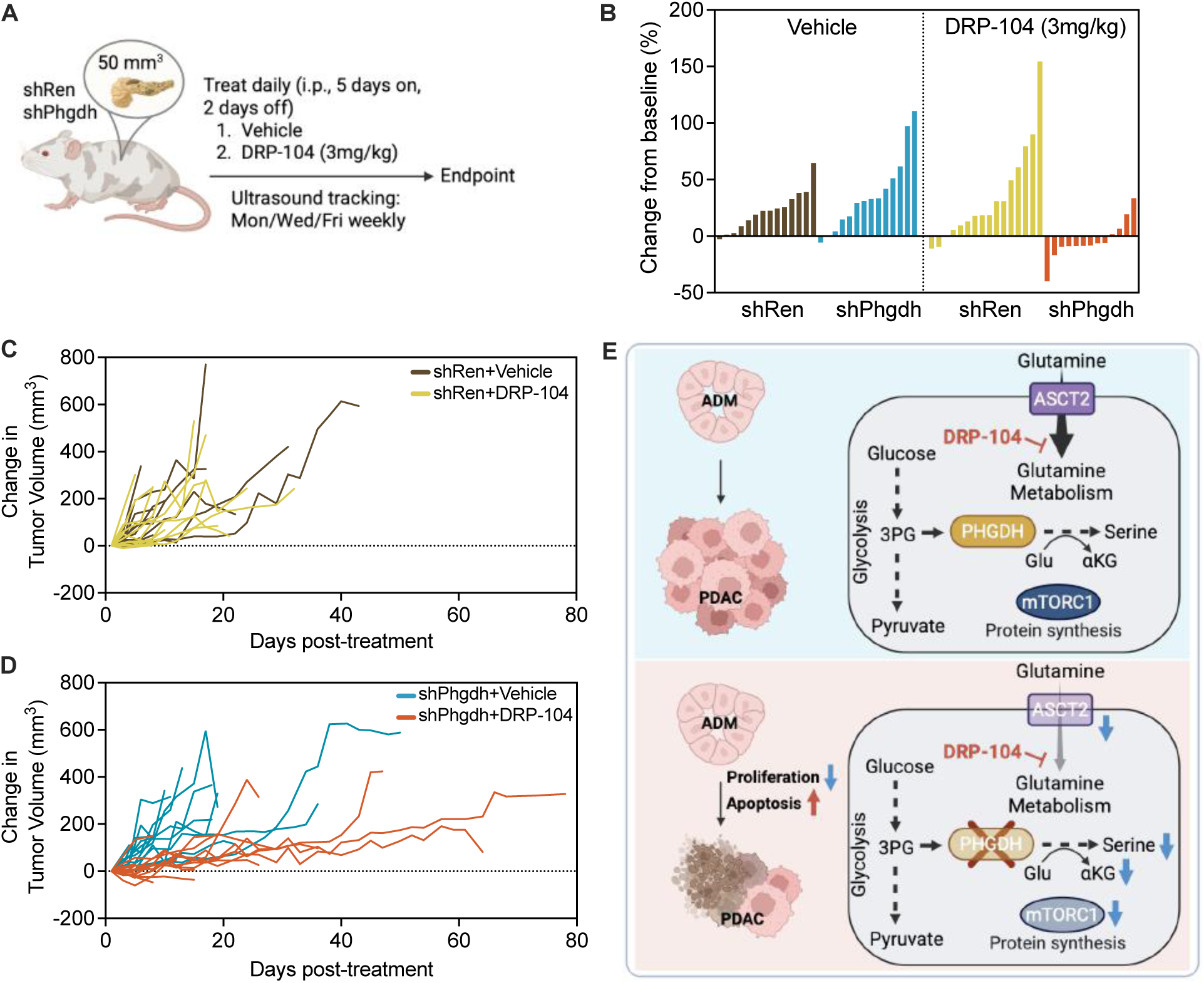
PHGDH deficiency sensitizes PDAC to glutamine antagonism. (A) Study schematic. shRen and shPhgdh PDAC mice were switched to doxycycline diet at postnatal day 30 and monitored for tumor development. Upon reaching ∼50 mm³ by ultrasound, mice were randomized to vehicle or DRP-104 (3 mg/kg, intraperitoneally, 5-days-on/2-days-off schedule). Tumor volume was measured by ultrasound three times per week, and tumors were collected at endpoint. (B) Waterfall plot showing change in tumor volume at the first on-treatment ultrasound assessment, normalized to baseline tumor volume at treatment initiation. shRen mice (vehicle, n = 14; DRP-104, n = 14) and shPhgdh mice (vehicle, n = 16; DRP-104, n = 13). (C and D) Longitudinal tumor volume measurements by ultrasound from treatment initiation to endpoint in (C) shRen cohorts treated with vehicle (n = 16) or DRP-104 (n = 15) and (D) shPhgdh cohorts treated with vehicle (n = 15) or DRP-104 (n =16). Lines represent individual mice. (E) Proposed model summarizing PHGDH-dependent PDAC progression and response to glutamine antagonism. In control tumors, PHGDH supports proliferation and sustains glutamine metabolism, resulting in a weak response to glutamine antagonism. Following PHGDH depletion, reduced ASCT2 abundance, diminished glutamine uptake, and reduced levels of glutamine-derived metabolites promote apoptosis and increase sensitivity to glutamine antagonism.

At the first on-treatment ultrasound, vehicle-treated shRen and shPhgdh tumors showed comparable growth (Fig. 4B). DRP-104-treated shRen tumors did not shrink and continued to increase in size, similar to vehicle-treated controls. In contrast, DRP-104 induced tumor regressions specifically in shPhgdh mice, with most mice showing decreases relative to baseline. Tumor volumes monitored over time showed that DRP-104 did not alter tumor growth compared with vehicle in shRen mice (Fig. 4C). In contrast, DRP-104 reduced tumor burden in shPhgdh mice relative to vehicle-treated controls, with tumors remaining smaller for much of the treatment period; however, a subset of tumors exhibited late outgrowth near endpoint (Fig. 4D). While overall survival from treatment initiation did not reach statistical significance across groups (Supplementary Fig. 3), few mice in the shPhgdh cohort treated with DRP-104 exhibited extended survival, consistent with the tumor responses observed by ultrasound. Together, these results demonstrate that PHGDH deficiency sensitizes PDAC tumors to glutamine antagonism.

## Discussion

PDAC develops within an extremely nutrient-deprived microenvironment yet sustains aggressive growth through distinctive metabolic adaptations, including atypical regulation of amino acid metabolism (4, 40, 41). We identified PHGDH as a stage-specific requirement for PDAC progression that operates through an unexpected mechanism. PHGDH suppression delayed malignant progression and induced apoptosis in established tumors. However, PHGDH-deficient PDAC did not compensate by increasing serine uptake as seen in other tumor types. Instead, PHGDH loss impaired glutamine uptake and metabolism, reduced ASCT2 expression, and created selective sensitivity to glutamine antagonism (Fig. 4E), revealing an unanticipated metabolic coupling between serine synthesis and glutamine handling.

Prior work showed that Kras-driven pancreatic cancer is largely insensitive to dietary serine and glycine restriction and proposed that SSP upregulation may preserve serine availability in PDAC (14). Studies in other tumor contexts suggest that impairment of endogenous serine synthesis can instead promote dependence on extracellular serine. For example, in colorectal cancer models, PHGDH inhibition increases reliance on exogenous serine and sensitizes tumors to serine limitation (36). We therefore tested whether PDAC tumors similarly compensate for impaired *de novo* serine synthesis by increasing uptake and utilization of exogenous serine. However, PHGDH-deficient PDAC tumors did not increase serine scavenging, and remained insensitive to extracellular serine and glycine availability.

The failure to engage compensatory serine uptake distinguishes PDAC from other tumor types and may be explained by the concurrent downregulation of ASCT2. As the primary transporter for serine and several other amino acids including glutamine, alanine, and threonine (42), reduced ASCT2 expression would impose a dual metabolic constraint: diminished *de novo* synthesis coupled with impaired capacity for extracellular amino acid acquisition. Importantly, in our model, PHGDH loss was restricted to tumor cells while host tissues retained intact serine biosynthesis, maintaining systemic serine availability. This differs from pharmacologic PHGDH inhibition studies in colorectal cancer, where systemic inhibitor treatment would be expected to reduce both tumor and host serine production, potentially limiting extracellular serine pools and amplifying dependence on dietary sources. Despite preserved host serine synthesis in our model, PDAC tumors failed to compensate through increased uptake. This mechanism likely extends beyond dietary sources to limit access to all extracellular serine pools. Notably, neuronal serine release was previously identified as an important mechanism supporting human PDAC growth in the setting of PHGDH deficiency (18). Additional work is needed to understand whether these tumors can maintain ASCT2 expression or upregulate alternative serine transporters (43) to compensate for ASCT2 loss.

The most striking metabolic consequence of PHGDH loss was impaired glutamine metabolism, an unanticipated phenotype that reveals a previously unrecognized link between serine biosynthesis and glutamine handling in PDAC. This coupling may reflect the coordinated requirement for both amino acids across multiple biosynthetic pathways. *De novo* purine synthesis requires serine-derived one-carbon units through the folate cycle as well as glutamine as a nitrogen donor at multiple steps. Similarly, pyrimidine and glutathione biosynthesis depend on both serine-derived intermediates and glutamine. The SSP itself consumes glutamate through PSAT1-catalyzed transamination. When PHGDH loss creates a serine-limited state, the cell’s capacity to productively utilize glutamine in these coupled biosynthetic pathways may become constrained. Maintaining high glutamine uptake without sufficient serine to support coordinated biosynthesis may be metabolically futile, providing a rationale for adaptive downregulation of glutamine acquisition (42). This positions PHGDH not simply as a serine biosynthetic enzyme, but as a regulator of broader biosynthetic capacity that coordinates serine and glutamine utilization.

The reduction in ASCT2 protein despite elevated mRNA suggests post-transcriptional regulation, potentially mediated through attenuated mTORC1 signaling, as evidenced by reduced p-S6. This is consistent with established mechanisms linking amino acid availability, mTORC1 activity, and transporter expression (44, 45). Whether PHGDH loss suppresses mTORC1 through direct effects on amino acid sensing pathways, energy stress, or secondary metabolic consequences remains to be determined. Importantly, ASCT2 functions as a bidirectional antiporter for multiple amino acids including glutamine, serine, alanine, and threonine (40), suggesting that its downregulation would impose broad constraints on amino acid metabolism beyond glutamine alone. This may explain both the failure to compensate through serine scavenging and the selective vulnerability to glutamine antagonism.

The synthetic lethality between PHGDH deficiency and glutamine antagonism has important therapeutic implications. Previous studies demonstrated that glutamine antagonism can suppress PDAC growth in preclinical models (46, 47), but clinical development has been limited by concerns about systemic toxicity and modest efficacy in unselected populations. Our findings suggest that PHGDH expression status may stratify tumors based on glutamine dependency, potentially identifying a subset of patients most likely to benefit from glutamine-targeted therapies. Moreover, the differential sensitivity to DRP-104 between PHGDH-high and PHGDH-low tumors raises the possibility of a therapeutic window that could be exploited through combined SSP and glutamine inhibition.

Several limitations of this study warrant consideration. First, our use of shRNA-mediated PHGDH knockdown in an ESC-GEMM model may not fully recapitulate the effects of pharmacologic PHGDH inhibition, which would produce systemic enzyme inhibition affecting both tumor and host tissues. While our acute pharmacologic studies with BI-9593 phenocopied key features of genetic PHGDH suppression, the long-term consequences of sustained systemic PHGDH inhibition on tumor evolution and host physiology remain to be determined. Second, the timing of doxycycline induction and PHGDH knockdown occurs after initial oncogenic transformation, potentially missing even earlier developmental requirements during the initiation of ADM from normal acinar tissue. Third, our KRAS; shp53-driven model represents one genetic context of PDAC, and it remains unclear whether these metabolic dependencies extend to tumors with different genetic alterations. Fourth, the mechanistic link between PHGDH loss and ASCT2 downregulation remains incompletely defined. While reduced mTORC1 signaling provides one plausible mechanism for decreased ASCT2 translation, the proximal signals connecting serine biosynthesis to mTORC1 activity require further investigation.

## Methods

### Mouse Models

All experiments were approved by the University of South Florida IACUC (protocols IS00012540R and IS00011831R). KPC PDAC mice (LSL-Kras^G12D/+^; Trp53^flox/flox^; p48-Cre) and KC PanIN mice (LSL-Kras^G12D/+^; p48-Cre) were generated by standard intercrossing. LSL-Kras^G12D/+^ [RRID:IMSR_JAX:008179], Trp53^flox^ [RRID:IMSR_JAX:008462], p48-Cre [RRID:IMSR_JAX:023329]. For inducible gene silencing studies, chimeric PDAC cohorts were generated using a rapid ESC-GEMM platform as described (35). Embryonic stem cells harboring LSL-KrasG12D, p48-Cre, CHC, and ROSA26-CAGs-LSL-rtTA3 alleles were obtained from the laboratory of Dr. Scott Lowe and engineered by RMCE to integrate TRE-driven GFP-miRE shRNA cassettes (shRen/shp53 or shPhgdh/shp53) into the Col1a1 locus. Correctly targeted clones were screened by PCR and injected into blastocysts to generate chimeric mice. shRNA expression was induced with doxycycline-containing diet (200ppm, AIN-93G diet, Teklad TD.94045) at 30 days of age.

### Histology and immunostaining

Pancreatic tissues were fixed in 10% formalin, paraffin-embedded, and sectioned at 4 μm thickness. H&E-stained sections were scanned using Aperio ScanScope AT2 (Leica Biosystems) at 20X magnification and analyzed in QuPath (v0.3.2) for disease and PDAC area quantification based on established morphologic criteria (48, 49). For immunohistochemistry, sections were deparaffinized in xylene and rehydrated through a graded ethanol series. Heat-mediated antigen retrieval was performed by boiling sections in 10 mM citrate buffer (pH 6.0). Endogenous peroxidase activity was quenched with 3% hydrogen peroxide, followed by blocking in 2.5% normal goat serum for 1 hours. Sections were incubated overnight at 4°C with primary antibodies against cleaved caspase-3 (Cell Signaling, #9664, 1:1000), Ki67 (Cell Signaling, #12202, 1:1000), PHGDH (Sigma, HPA021241, 1:1000), GFP (Cell Signaling, #2956, 1:1000), phospho-S6 [S240/244] (Cell Signaling #5364, 1:1000), and ASCT2 (Thermo 20350-1-AP, 1:100). Immunoreactivity was detected using an HRP polymer detection system according to the manufacturer’s instructions (ImmPRESS HRP kit, Vector Laboratories, RRID: AB_2631198, matched to the host species of the primary antibody). Signal was developed using DAB substrate (Vector Laboratories, SK-4105) and sections were counterstained with hematoxylin (Vector Laboratories, H-3404), dehydrated, and mounted. For immunofluorescence, sections were permeabilized, blocked, and incubated with primary antibodies against GFP (Cell Signaling, #2956, 1:200) and phospho-S6 [S240/244] (Cell Signaling, #5364, 1:200). After washing, sections were incubated with Alexa Fluor-conjugated secondary antibodies: donkey anti-rabbit IgG (H+L), Alexa Fluor 488 (Invitrogen, A-21206) for phospho-S6 and goat anti-mouse IgG (H+L), highly cross-adsorbed, Alexa Fluor 647 (Invitrogen, A-21236) for GFP. Sections were mounted using a mounting medium containing DAPI (Vector Laboratories, H-1200). Quantification was performed in QuPath using H-score and percent-positive cells or mean fluorescence intensity.

### RNAscope In Situ Hybridization

RNAscope 2.5 chromogenic in situ hybridization was performed on FFPE PDAC sections to detect Slc1a5 mRNA using the RNAscope 2.5 assay (ACD RNAscope® 2.5 HD Detection Reagent - BROWN, 322310) following the manufacturer’s instructions. After hybridization with the Slc1a5 probe (RNAscope™ Probe – Mm-Slc1a5, #455201) and signal amplification, slides were counterstained with hematoxylin, dehydrated through graded ethanol, cleared in xylene, and mounted with (Vector Laboratories, Cat# H-5700-60). RNAscope puncta were quantified in QuPath (v0.3.2) using an analysis workflow guided by the RNAscope QuPath technical note (ACD Bio-Techne). After hematoxylin-based cell segmentation, PDAC cells were manually curated by excluding non-tumor cells. Subcellular detection was used to quantify DAB puncta with empirically optimized detection parameters to minimize background. Each cell was assigned an RNAscope score (0–4) using established criteria, and H-score (%) was calculated as: 100 × (N1 × 1 + N2 × 2 + N3 × 3 + N4 × 4) / Ntotal, where N1–N4 are the numbers of cells assigned RNAscope scores 1 to 4 and Ntotal is the total number of cells analyzed. Representative images were captured using the Axio Lab A.1 microscope (Carl Zeiss Microimaging Inc.) at 100X magnification.

### Stable isotope tracing

In vivo isotope tracing was performed using continuous tracer infusion in awake animals. Depending on the tracer and route, mice were catheterized via either the jugular vein or tail vein using our established procedures (50). Jugular vein catheters were surgically implanted 3–7 days prior to infusion to allow recovery from surgery, whereas tail vein catheters were placed on the day of infusion. During infusions, mice were housed in a mouse harness with tether and swivel (SAI Infusion Technologies) to prevent catheter dislodgement while allowing free movement within the cage. Tracer solutions were prepared fresh in sterile saline, passed through a 0.22 µm filter, and infused using a syringe pump (BS-8000, Braintree Scientific). U-^13^C_6_-glucose (Cambridge Isotope Laboratories, CLM-1396, 150mg/mL) was infused via the jugular vein at 100 µL/min for 1 minute, followed by a continuous infusion at 3 µL/min for 2.5 hours. ^13^C_1_-serine (Cambridge Isotope Laboratories, CLM-1573, 20mg/mL) was infused via the jugular vein at 2 µL/min for 4 hours. ^13^C_5_-glutamine (Cambridge Isotope Laboratories, CLM-1822, 15 mg/mL) was infused via the tail vein at 2 µL/min for 4 hours.

At the end of each infusion, blood was collected via the submandibular vein into serum separator tubes (BD, cat# 365967) and kept on ice until serum isolation. Mice were euthanized by cervical dislocation, and tumors and indicated tissues were rapidly dissected and snap-frozen in liquid nitrogen. Serum and tissues were stored at −80°C until LC-HRMS analysis.

### LC-MS analysis and data processing

Frozen tumors were pulverized under cryogenic conditions using a pre-chilled tissue pulverizer (BioSpec, 59012MS). Metabolites were extracted by adding ice-cold 80% methanol (pre-chilled to −80°C) at a final concentration of 50 mg tissue per mL of extraction solvent, followed by incubation at −80°C for 24 hours. For endpoint tumor metabolomics comparing shRen and shPhgdh samples, tumor samples were extracted using an 80% methanol-based solvent containing 5 mM N-ethylmaleimide (NEM) prepared in 10 mM ammonium formate (pH 7.0), and extracts were stored at -80°C overnight prior to centrifugation. For serum extraction, 10 µL of serum was mixed with 90 µL of ice-cold extraction solvent (88.8% methanol, −80°C), vortexed for 10 seconds, and incubated at −80°C for 15 minutes. Extracts were clarified by centrifugation at 17,000 g for 20 minutes at 4°C, and supernatants were analyzed by LC-HRMS on a Vanquish UHPLC system coupled to a Q Exactive HF Orbitrap mass spectrometer equipped with a heated electrospray ionization source (Thermo Fisher Scientific). Metabolites were separated using an Atlantis Premier BEH Z HILIC VanGuard FIT column (2.1 × 150 mm, 2.5 µm; Waters) maintained at 30°C. Mobile phase A consisted of 10 mM ammonium carbonate with 0.05% ammonium hydroxide in water, and mobile phase B consisted of acetonitrile. The gradient was 80% B at 0 min, ramped to 20% B at 13 min, held at 20% B until 15 min, and then returned to initial conditions for re-equilibration. The flow rate was 150 µL/min and the injection volume was 5 µL. MS data were acquired in positive electrospray ionization mode. Full-scan MS1 spectra were collected over m/z 65–950 at a resolution of 120,000 (at m/z 200) with an AGC target of 3 × 10^6^. Raw files were converted to .cdf format using Xcalibur (v4.0) and processed in El-Maven (v0.12.0). Peaks were extracted using an EIC mass tolerance of ±10 ppm, and metabolite assignments were made based on accurate mass and retention time matching to an in-house library of authentic standards. Peak areas for labeled and unlabeled isotopologues were integrated using El-Maven (v0.6.1) or Thermo Xcalibur Qual Browser. Isotopologue abundances were corrected for naturally occurring isotope abundance using IsoCor (v2.2.0).

### Dietary and Pharmacological Interventions

For serine/glycine studies, mice received (i) control diet (Envigo, TD.110839) plus control drinking water, (ii) serine and glycine-free diet (Envigo, TD.160752) plus control drinking water, or (iii) control diet (Envigo, TD.110839) plus 2% (w/v) serine and glycine supplementation in drinking water. All diets contained 200ppm doxycycline. Diets were changed weekly and drinking water was changed twice weekly. BI-9593 or inactive control BI-9594 were formulated in 0.5% Natrosol administered by oral gavage at the indicated frequency. DRP-104 was administered at 3 mg/kg in Tween80:ethanol:saline (5:5:90, v/v/v) by intraperitoneal injection on a 5-days-on/2-days-off schedule as previously described (46).

### Ultrasound imaging

Ultrasound imaging was performed using the Vevo F2 system (FUJIFILM VisualSonics) at the SAIL core facility. Mice were anesthetized with isoflurane in oxygen according to the facility standard protocol and maintained on a heated imaging platform to prevent hypothermia. Abdominal hair was removed using clippers and depilatory cream, and ultrasound gel was applied prior to image acquisition. Tumor imaging was performed in B-mode with 3D acquisitions using the UHF57x transducer, and tumor volumes were quantified using Vevo Lab software (v5.7.1) with the facility’s standard analysis workflow. Ultrasound imaging was performed once weekly prior to treatment initiation and then three times per week during DRP-104 treatment until endpoint.

### Statistical analysis

Statistical analyses were performed using GraphPad Prism (v9). Two-group comparisons were assessed using the Mann-Whitney test. Survival curves were compared using the log-rank (Mantel–Cox) test. Data are presented as mean ± SD unless otherwise indicated, and individual mice are shown as dots when displayed. P-values < 0.05 were considered statistically significant.

## Author Contribution

Y.K. and G.M.D. conceived the study, designed the experiments, and wrote the manuscript. Y.K. performed the experiments and analyzed and interpreted the data. L.J.S. contributed to histopathologic annotations. S.C. and M.L. assisted with animal experiments. H.C.M. and K.P.O. contributed human PanIN and PDAC specimen data and related analyses. F.K. contributed to the development of the PHGDH-targeting ESC-GEMM model. G.M.D. supervised the project, acquired funding, and edited the manuscript. All authors discussed the results and commented on the manuscript.

## Acknowledgments

We thank members of the DeNicola laboratory for helpful discussions. We also thank members of the Karreth and Gomes laboratories for sharing reagents and for helpful input throughout the study. We are grateful to Dan K. Lester and Joseph L. Kissil for providing RNAscope reagents and technical guidance. We also thank Joseph Johnson for microscopy support and helpful advice on image acquisition and analysis, as well as William Dominguez Viqueira, Filip Konecny, Epi Ruiz, and Alex J. Lundberg for assistance with ultrasound imaging.

This work was supported by the PanCAN/AACR Pathway to Leadership Award (G.M.D.), NIH/NCI grants R21CA289213 (F.A.K.) and R01CA215607 (K.P.O.), and the 2025 Miles for Moffitt–Team Science Postdoctoral Award (Y.K.). This work was also supported by the Gene Targeting Core, Analytic Microscopy Core, Small Animal Imaging Lab Core, and Proteomics and Metabolomics Core Facilities at the H. Lee Moffitt Cancer Center & Research Institute, which are supported in part by the Moffitt Cancer Center Support Grant (P30CA076292). All schematics were created with BioRender.com.

## Supplementary Information

**Supplementary Figure 1.**
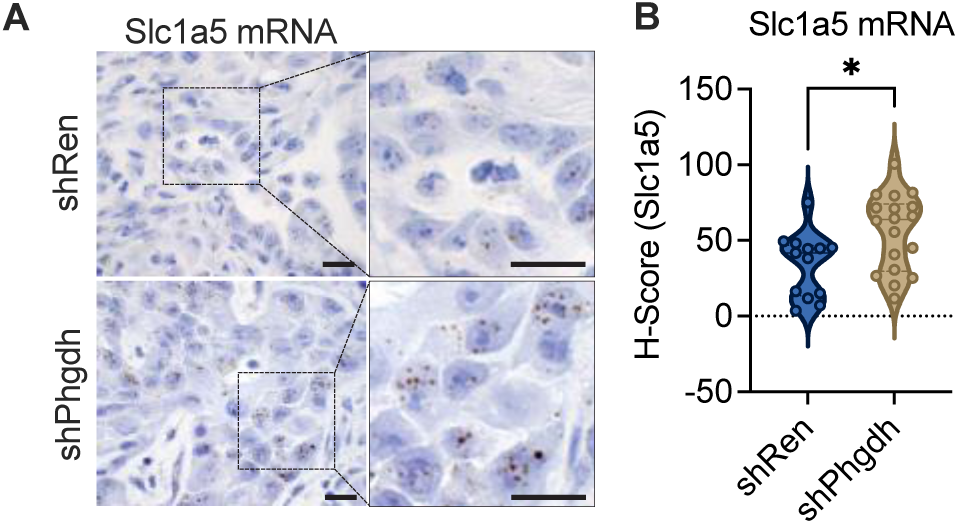
PHGDH silencing does not decrease Slc1a5 mRNA. (A) Representative RNAscope 2.5 HD-BROWN (DAB) images for Slc1a5 mRNA in endpoint PDAC tumors from shRen and shPhgdh mice. Left, images acquired using a 100x objective; right, enlarged views of the boxed regions. Scale bars = 10 µm. (B) H-score quantification of Slc1a5 RNAscope signal in endpoint PDAC tumors (shRen, n = 13; shPhgdh, n = 18).

**Supplementary Figure 2.**
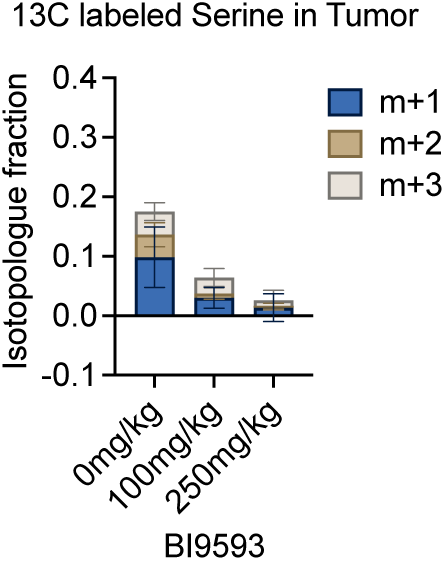
Does-dependent inhibition of Phgdh with BI-9593 in vivo. Endpoint shREN mice (41–67 days post-doxycycline) received BI-9593 by oral gavage at 0, 25, 100, or 250 mg/kg (n = 3/group). At 4.5 hours post-treatment, mice were infused with U-^13^C_6_-glucose for 2.5 h, and tumors were collected for metabolomic analysis. Serine isotopologue fractions (M+1, M+2, M+3) in tumors were quantified by LC-MS. Bars show mean ± SD.

**Supplementary Figure 3.**
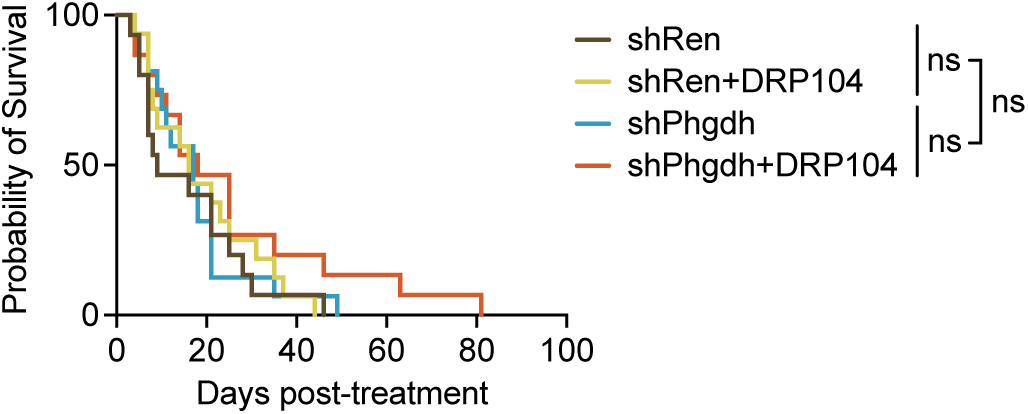
Overall survival following glutamine antagonism in shRen and shPhgdh PDAC mice. Kaplan–Meier overall survival from treatment initiation (defined as the day tumors reached ∼50 mm³ by ultrasound) in shRen and shPhgdh cohorts treated with vehicle or DRP-104 (3 mg/kg, i.p., 5 days on/2 days off). shRen+vehicle (n = 15), shRen+DRP-104 (n = 16), shPhgdh+vehicle (n = 16), and shPhgdh+DRP-104 (n = 15). Survival was monitored until endpoint. Statistical comparisons were performed using the log-rank (Mantel–Cox) test; ns, not significant.

**Supplementary Table 1. Differential metabolite abundance in endpoint PDAC tissues from shRen and shPhgdh mice.** The table includes normalized values, fold changes, and p-values from endpoint PDAC tissues of shRen and shPhgdh mice (n = 9 per group) used to generate the volcano plot in Figure 3A.

